# Stable hypermutators revealed by the genomic landscape of DNA repair genes among yeast species

**DOI:** 10.1101/2025.03.15.643480

**Authors:** Carla Gonçalves, Jacob L. Steenwyk, David C. Rinker, Dana A. Opulente, Abigail L. LaBella, Marie-Claire Harrison, John F. Wolters, Xiaofan Zhou, Xing-Xing Shen, Shay Covo, Marizeth Groenewald, Chris Todd Hittinger, Antonis Rokas

## Abstract

Mutator phenotypes are short-lived due to the rapid accumulation of deleterious mutations. Yet, recent observations reveal that certain fungi can undergo prolonged accelerated evolution after losing DNA repair genes. Here, we surveyed 1,154 yeast genomes representing nearly all known yeast species of the subphylum Saccharomycotina to examine the relationship between reduced DNA repair repertoires and elevated evolutionary rates. We identified three distantly related lineages—encompassing 12% of species—with substantially reduced sets of DNA repair genes and the highest evolutionary rates in the entire subphylum. Two of these “faster-evolving lineages” (FELs)—a subclade within the order Pichiales and the *Wickerhamiella*/*Starmerella* (W/S) clade (order Dipodascales)—are described here for the first time, while the third corresponds to a previously documented *Hanseniaspora* FEL. Examination of DNA repair gene repertoires revealed a set of genes predominantly absent in these three FELs, suggesting a potential role in the observed acceleration of evolutionary rates. Genomic signatures in the W/S clade are consistent with a substantial mutational burden, including pronounced A|T bias and signatures of endogenous DNA damage. The W/S clade appears to mitigate UV-induced damage through horizontal acquisition of a bacterial photolyase gene, underscoring how gene loss may be offset by nonvertical evolution. These findings highlight how the loss of DNA repair genes gave rise to hypermutators that persist across macroevolutionary timescales, with horizontal gene transfer as an avenue for partial functional compensation.

## Introduction

Mutations are the raw material of evolution and adaptation. Since mutations are more likely to be deleterious than adaptive (Eyre-Walker and Keightley, 2007), cells have evolved numerous interacting mechanisms to ensure high fidelity of genome replication and stability—such as cell cycle checkpoints and DNA damage sensing and repair pathways (Giglia-Mari, et al. 2011; Kreuzer 2013; Steenwyk 2021). Impairment in DNA repair pathways has been associated with hypermutation across the tree of life (Bridges 2001; Roberts and Gordenin 2014; Campbell, et al. 2017; Steenwyk, et al. 2019; Phillips, et al. 2021; Gambhir, et al. 2022). Hypermutation can arise from the aberrant function of genes involved in several pathways, such as DNA damage/S-phase checkpoints, cell cycle, DNA replication, or oxidative stress (Murakami-Sekimata, et al. 2010; Serero, et al. 2014). For example, defects in the mismatch repair (MMR) system, a highly conserved pathway that corrects mismatched bases produced during DNA replication (Fukui 2010), have been shown to lead to diverse mutator phenotypes (Oliver, et al. 2000; Chopra, et al. 2003; Reis, et al. 2019; Gambhir, et al. 2022). In bacteria, deficiency in the MMR can also result in the relaxation of recombination barriers between species, resulting in higher horizontal gene transfer (HGT) rates between distantly related species (Rayssiguier, et al. 1989; Thomas and Nielsen 2005).

Hypermutators have been mostly observed in fluctuating environments (Swings, et al. 2017; Callens, et al. 2023), such as in the context of human disease (Oliver, et al. 2000; Chopra, et al. 2003; Healey, et al. 2016; Billmyre, et al. 2017; Rhodes, et al. 2017; Gambhir, et al. 2022). In clinical isolates of bacterial and fungal pathogens (Healey, et al. 2016; Billmyre, et al. 2017; Rhodes, et al. 2017; Gambhir, et al. 2022) , defects in DNA repair pathways have been associated with increased mutation rates, which are thought to facilitate host adaptation. For example, mutations in the MMR gene *MSH2* in *Cryptococcus* (Billmyre, et al. 2017; Rhodes, et al. 2017) or *Nakaseomyces glabratus* (Healey, et al. 2016) opportunistic fungal pathogens are known to give rise to mutator phenotypes, possibly promoting rapid host adaptation and drug resistance.

Hypermutator phenotypes have been also implicated in the acceleration of disease progression and the evolution of resistance to treatments in humans (Campbell, et al. 2017; Jiang, et al. 2020).

Fluctuating environments can, therefore, serve as triggers for the emergence and maintenance of hypermutator phenotypes because they enable a broader exploration of genotype-phenotype space and access to a larger pool of potentially advantageous mutations (Shaver, et al. 2002). However, once these beneficial mutations are fixed in the population, and organisms become well adapted to their new environment, compensatory mutations that reduce the mutation rate will be favored (Kimura 2009; Wielgoss, et al. 2013). Genetic constraints, such as the loss of DNA repair genes, may, however, constrain the re-lowering of mutation rates.

In fungi, the DNA repair gene repertoire has been described to be highly variable (Milo, et al. 2019; Shen, et al. 2020), and in a few ancient lineages, loss of DNA repair-related genes has been associated with long-term increased evolutionary rates (Steenwyk, et al. 2019; Phillips, et al. 2021). Among these lineages include a case of macroevolutionary hypermutation among *Hanseniaspora* yeast (Steenwyk, et al. 2019). These examples suggest that hypermutator lineages can survive and diversify over macroevolutionary timescales. However, the prevalence of such long-term hypermutator lineages, the extent of their association with the loss of DNA repair genes, and the mechanisms involved in their long-term survival remain poorly understood.

Here, we used a sequence similarity search approach, partially validated by structural homology and phylogenetic approaches to explore the relationship between DNA repair gene repertoires and evolutionary rates across the entire subphylum Saccharomycotina, currently comprising more than 1,000 species (Groenewald, et al. 2023; Opulente, et al. 2024). We found that the three lineages with the highest rates of sequence evolution (faster-evolving lineages or FELs) in the subphylum have experienced substantial reductions in their DNA repair repertoires. These are a subclade within the order Pichiales, the *Wickerhamiella*/*Starmerella* (W/S) clade (order Dipodascales), and a previously reported *Hanseniaspora* lineage (order Saccharomycodales) (Steenwyk, et al. 2019). Several genes have been seemingly independently lost in the three FELs, suggesting a possible role in accelerating mutation rates. In the W/S clade, we found strong signatures of mutational burden compared to its closest relatives—namely, a pronounced A|T bias and increased frequency of mutations associated with endogenous DNA damage, but no evidence of UV damage. Interestingly, most W/S-clade species harbour either a filamentous fungal-like version or a bacterial version of *PHR1* - likely horizontally acquired and seemingly functional - while most W/S clade closest relatives lack *PHR1*. These results suggest that variation of evolutionary rates through the loss of DNA repair genes may have been common in Saccharomycotina yeast evolution and that other evolutionary mechanisms, such as HGT, might help circumvent constraints imposed by loss.

## Results

### Extensive variation in DNA repair gene repertoire across the Saccharomycotina

Recent investigations of gene family and trait evolution of the Saccharomycotina subphylum have shown that losses of genes and traits are frequent and substantially contribute to yeast diversity (Shen, et al. 2018; Feng, et al. 2024). For example, research in the genus *Hanseniaspora* revealed a substantial loss of DNA repair genes, which seems to be associated with rapid evolution in this lineage (Steenwyk, et al. 2019). However, it remains unclear how DNA repair repertoires and evolutionary rates vary across the entire yeast subphylum, we examined the distribution of 415 DNA repair-related proteins (henceforth referred to as DNA repair genes (Table S1) across 1,154 proteomes representing 1,090 species within the subphylum Saccharomycotina (Groenewald, et al. 2023; Opulente, et al. 2024) using Hidden Markov Model sequence similarity searches (Steenwyk, et al. 2019; Phillips, et al. 2021; Steenwyk and Rokas 2021).

We observed that DNA repair gene repertoire extensively varied across Saccharomycotina (Fig. 1A, Table S1, Table S2). Some genes were widely conserved, such as *RAD3* encoding a DNA helicase involved in nucleotide excision repair and transcription (Naumovski and Friedberg 1983), which was present in all but one strain. Other genes were poorly conserved, such as *IRC4*, which encodes a protein involved in double-strand break repair and was absent from 98% of the proteomes inspected. Of the 415 genes examined, 225 (54.22%) were found in all proteomes; in contrast, 108 (26.02%) were absent in > 10 proteomes (Table S1). Genes considered to be essential in *S. cerevisiae* were generally highly conserved across Saccharomycotina (Table S1), but we also found several exceptions. For instance, *ABF1*, encoding a transcription factor involved in chromatin silencing and remodeling (Rhode, et al. 1989; Rhode, et al. 1992), was absent in more than 80% of the proteomes examined (Table S1).

**Figure 1.**
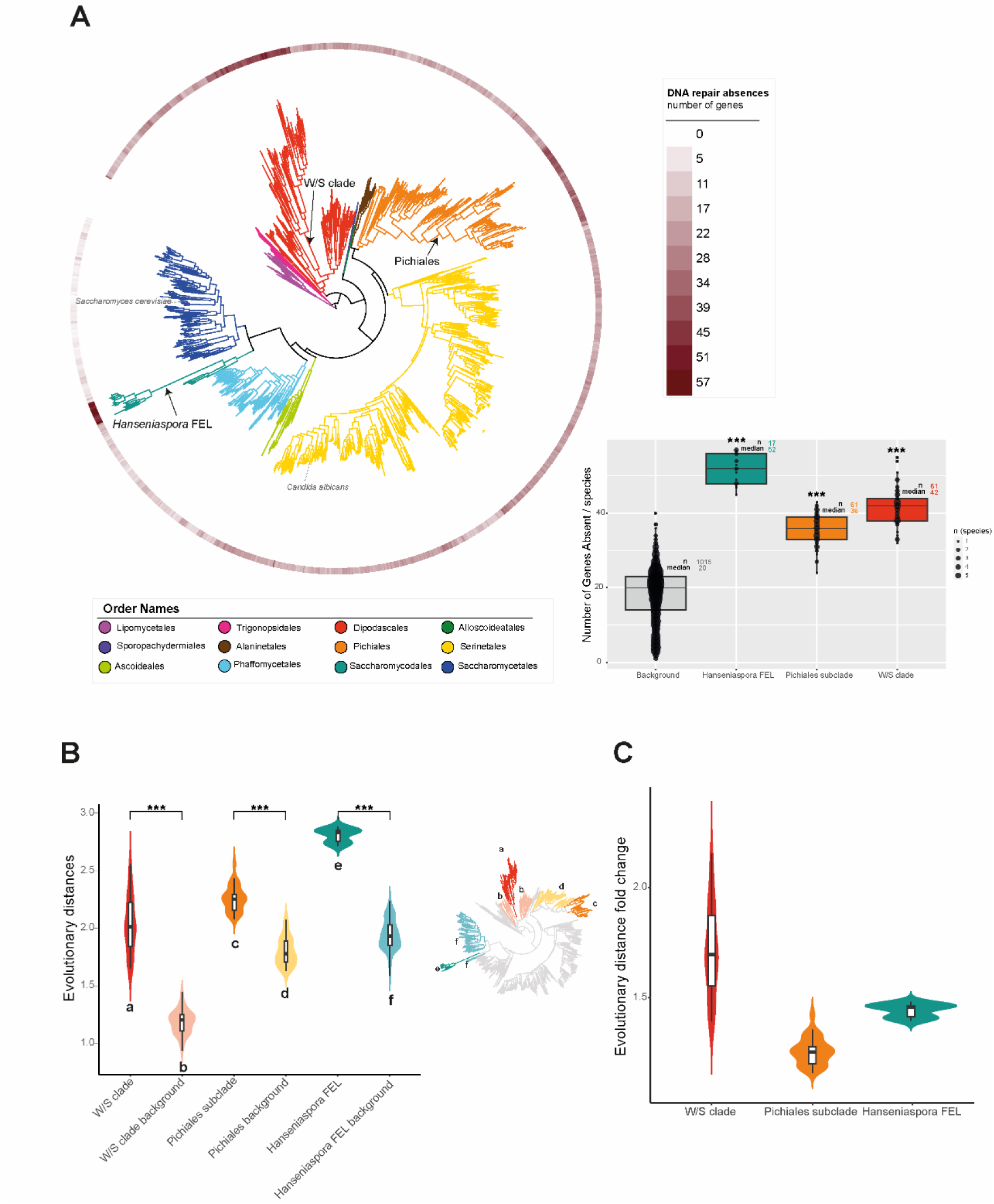
The DNA repair gene repertoire is highly variable across the subphylum Saccharomycotina and significantly reduced in three faster-evolving lineages. A) Distribution of the number of absent DNA repair genes (from a total of 415) across the Saccharomycotina species phylogeny depicting three distantly related lineages that lack the largest number of DNA repair genes: W/S clade, Pichiales subclade, and *Hanseniaspora* FEL. The panel on the right shows the number of absent genes across these three lineages (foreground species) compared to the number of genes absent in the background species (all others). B) Evolutionary distances were determined using tip-to-root distances of the Saccharomycotina species phylogeny from Opulente et al., 2024 (Opulente, et al. 2024) as proxy. C) Fold change difference between tip-to-root distances in the foreground species of each foreground lineage in respect to the average tip-to-root distance in the closest relatives (background). Statistical significance (panels A and B) was assessed using a pairwise Wilcoxon rank test after testing for normality. P-values were adjusted using the Holm correction (*** p-value < 0.001). In panel B, statistical significance was assessed between each of the faster-evolving lineages and the background. The Saccharomycotina species phylogeny was obtained from Opulente et al. 2024 (Opulente, et al. 2024). For reference, phylogenetic position of *Saccharomyces cerevisiae* and *Candida albicans* are shown in the tree.

Species belonging to Saccharomycetales had the largest DNA repair gene repertoire; other lineages generally had smaller DNA repair gene repertoires (Table S1, Fig. 1A). The lineage with the smallest repertoire was the *Hanseniaspora* FEL (Fig. 1A, Table S2), which was previously reported to have lost multiple genes involved in DNA repair and cell cycle pathways (Table S2) (Steenwyk, et al. 2019). Two additional lineages were identified as having substantially reduced DNA repair gene repertoires compared to their closest relatives and other Saccharomycotina species (Fig. 1A, Fig. S1, Table S2). One lineage comprised species belonging to the genera *Pichia*, *Saturnispora*, and *Martiniozyma*, all in the order Pichiales (henceforth referred to as the Pichiales subclade). Within this subclade, the average number of absent DNA repair genes (36) exceeds the average for Pichiales as a whole (approximately 19) (Fig. 1B). The second clade includes species belonging to the genera *Wickerhamiella* and *Starmerella* (W/S clade) (Fig. 1A). In the W/S clade, the number of genes absent ranges from 32-55, while the number of missing genes ranges from 16 to 29 in close relatives belonging to the genera *Blastobotrys* and *Sugiyamaella*. Notably, the three lineages—*Hanseniaspora,* Pichiales subclade, and the W/S clade—are all distantly related, suggesting that the reductions of their DNA repair gene repertoires have taken place independently.

### Independent DNA repair gene losses are associated with accelerated evolutionary rates

We next determined if evolutionary rate variation is associated with DNA repair gene repertoire. The species with the highest evolutionary rates belonged to the three lineages with the higher proportions of DNA repair absences: *Hanseniaspora* FEL, the Pichiales subclade, and the W/S clade (p-value < 0.001, Wilcoxon rank test, Fig. 1A, Fig. S2A). Evolutionary rates among FELs are significantly higher (p-value < 0.001, Wilcoxon rank test) compared to their respective closest relatives (Fig. 1B, Fig. 1C). The differences in evolutionary rates between these clades and their closest relatives (evolutionary rate fold change), were pronounced in the W/S clade and in *Hanseniaspora* FEL (Fig. 1C). Evolutionary rates were relatively uniform within *Hanseniaspora* FEL and in the Pichiales subclade (2.71–2.88 substitutions per site in *Hanseniaspora* FEL and 2.08–2.58 substitutions per site in the Pichiales subclade) but highly variable in the W/S clade (1.65 – 2.56 substitutions per site; Fig. 1B), which can be partially attributed to the fact that *Starmerella* species exhibit slightly higher evolutionary rates than *Wickerhamiella* species (Fig. S2A).

In *Hanseniaspora* FEL, the elevated evolutionary rates were previously found to be concentrated on the stem branch leading to the clade (Steenwyk, et al. 2019). Interestingly, terminal branches appeared to contribute substantially to the observed rate differences in both the W/S clade and the Pichiales subclade (Fig. 1A). We next tested whether there was a correlation between the total number of absent DNA repair genes and evolutionary rate across the yeast phylogeny using a phylogenetically corrected analysis. We found that the correlation is statistically significant (p-value = 4.5e^-10^, Phylogenetically Independent Contrasts – PIC method, Fig. S2B) but weak (adjusted R-squared: 0.04074), suggesting that there is no consistent link between the size of the DNA repair gene repertoires and altered evolutionary rates.

### Some DNA repair genes are predominantly absent in the faster-evolving lineages

Genes consistently absent across all three FELs may represent strong candidates for contributing to accelerated evolutionary rates (Fig. S3). To evaluate this, we first selected genes absent in more than 30% of strains of at least two foreground lineages (Fig. S4) and compared these genes among all foreground strains (*Hanseniaspora* FEL, W/S clade and Pichiales subclade, N=139) to the background strains (all others, N=1,015) (Fig. S4). We found that 17 of the 24 (70.83%) selected genes are also absent in a high proportion of background species (Fig. S4), suggesting that a substantial fraction of the losses is not specific to the foreground clades.

Seven genes (*MHF2*, *ISY1*, *MAG1*, *MGT1*, *SLX4*, *MTE1*, and *AHC1*) were found to be absent in more than 50% of all foreground strains and only in less than 10% of their background counterparts (Fig. 2B, Fig. S4). These genes are involved in multiple pathways in *S. cerevisiae*. For instance, *MHF2* is part of the MHF histone-fold complex, along with *MHF1*, which is involved in cellular response to DNA damage stimulus (Yang, et al. 2012). We found that *MFH1* was in more than 50% of the FEL strains lacking *MHF2*, but these proteins are only 90 amino acids long which might hinder their accurate identification. *ISY1* is part of the NineTeen complex and is involved in the regulation of the fidelity of pre-mRNA splicing (Villa and Guthrie 2005); *MAG1* is involved in the base excision repair (BER) pathway; *MGT1* contributes to repairing DNA alkylation (Xiao, et al. 1991); *MTE1* is involved in maintenance of telomere length and double-strand break repair (Yimit, et al. 2016); and *AHC1* is a subunit of the Ada histone acetyltransferase complex involved in double strand break repair (Eberharter, et al. 1999). *SLX4* encodes one of the subunits of the Slx1-Slx4 endonuclease involved in double-strand break repair (Coulon, et al. 2004; Pardo and Aguilera 2012; Gaur, et al. 2015; Covo 2020). In line with this, we found that all predicted W/S proteomes lacking Slx4 also lack Slx1, while *Hanseniaspora* FEL strains that lack Slx4 contain Slx1.

**Figure 2.**
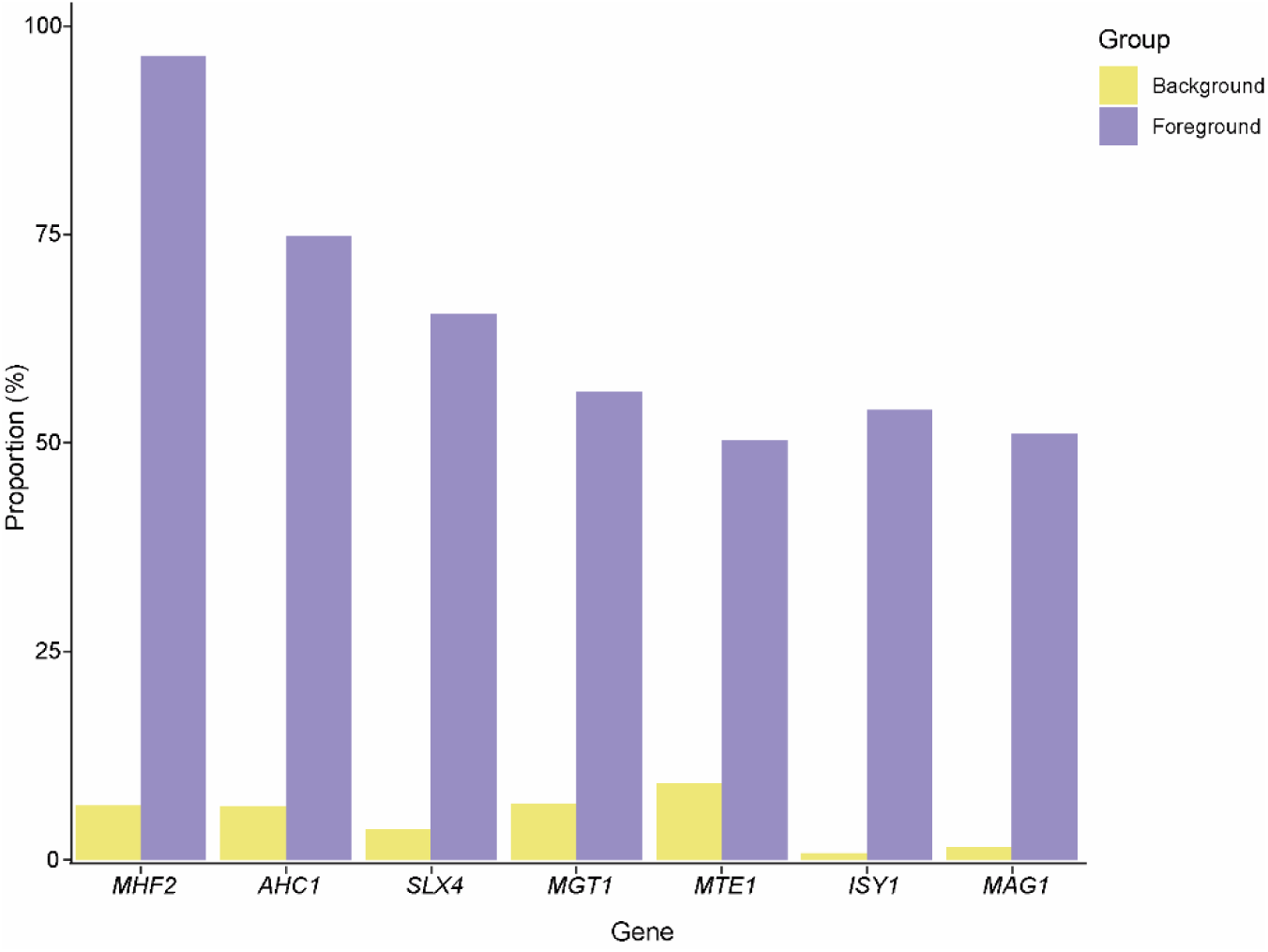
Faster-evolving lineages show overlapping absences of seven DNA repair genes. Genes absent in more than 50% of the total foreground strains (W/S clade + *Hanseniaspora* FEL + Pichiales subclade, N=139; shown in purple bars) and in less than 10% in the background species (N=1,015; shown in yellow bars) are represented. The Y axis represents the proportion of species in which each gene is absent is represented in foreground and background groups. This proportion was calculated based on the number of proteomes in which each gene was absent out of the total number of proteomes inspected.

After confirming the distribution of these genes through tBLASTn against the genomes and validating presences for proteins with a patchy distribution in all three clades, we found that some lineages almost or completely lack certain genes. For instance, *MAG1* is absent in all *Hanseniaspora* FEL strains inspected, while *MGT1* is absent in 70% of the strains, in line with previous reports (Steenwyk, et al. 2019). *MAG1* and *MGT1* are also inferred to be absent from all strains in the Pichiales subclade, except for *Candida sorboxylosa*, which contains an *MGT1* gene. A BLASTp search against the NCBI nr database revealed that the *MGT1* homologs present in both *C. sorboxylosa* and 30% of the *Hanseniaspora* FEL are likely of bacterial origin suggesting that they were acquired through HGT (Fig. S5). However, topology tests could not reject the null hypothesis of vertical descent (p-value > 0.05, Approximately Unbiased [AU] test). In the W/S clade, all strains lack *AHC1,* and most species lack *ISY1. AHC1* is also absent in all Pichiales genomes inspected and *ISY1* is absent in almost all *Hanseniaspora* FEL.

While not all genes are directly involved in DNA repair (e.g., *ISY1*), others, such as *MAG1,* are and might therefore be involved in increasing mutation rates. Furthermore, the independent losses of these genes and respective acceleration of mutation rates likely occurred at different evolutionary time points. Relaxed molecular clock analyses (Opulente et al., 2024) estimate that the onset of accelerated mutation rates occurred approximately 87 million years ago (mya) in the *Hanseniaspora* FEL and 73 mya in the Pichiales subclade, coinciding with their divergence from their closest relatives. In the W/S clade, this acceleration appears to have originated substantially earlier, around 253 mya.

### Reduction in DNA repair gene repertoire not always associated with accelerated evolutionary rates

In all the three faster-evolving lineages inspected, we observed that high evolutionary rates were associated with a substantial loss of DNA repair genes. To further investigate the relationship between DNA repair gene loss and evolutionary rate acceleration, we decided to focus on the W/S clade, which comprises floral species belonging to the *Starmerella* and *Wickerhamiella* genera (Lachance, et al. 2000; Lachance, et al. 2001; Gonçalves, et al. 2020), because the differences in evolutionary rates between this clade and their closest relatives were particularly pronounced (Fig. 1). Furthermore, the W/S-clade sister lineage, which contains only three species [*Candida incommunis*, *Candida bentonensis*, and *Deakozyma indianensis* (Groenewald, et al. 2023; Opulente, et al. 2024)], showed no evidence of acceleration of evolutionary rates (Fig. 3B) even though it too experienced numerous DNA repair gene losses (between 33-36 genes were absent, Fig. 3A).

**Figure 3.**
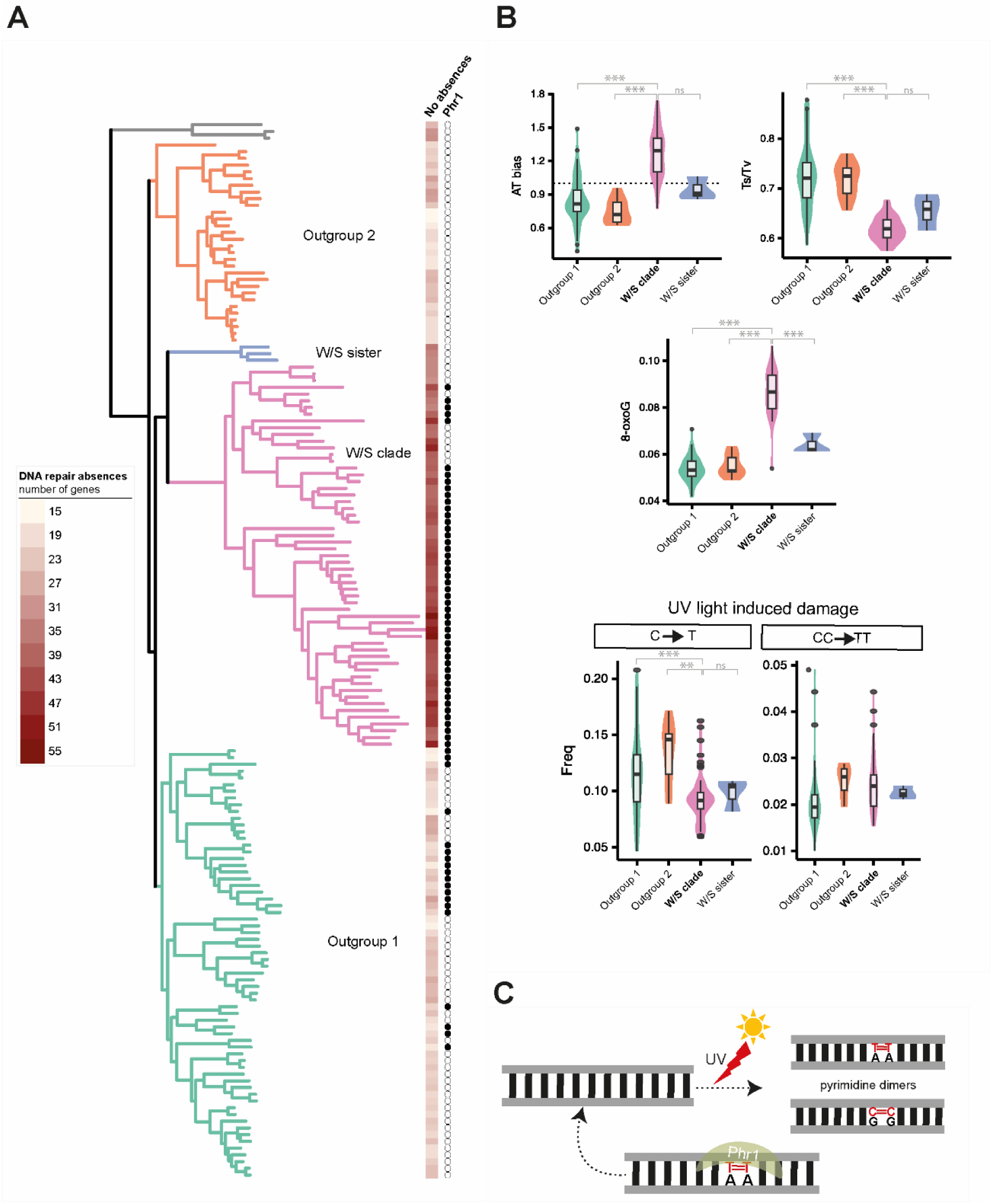
Absence of DNA repair genes in the W/S clade is commensurate with mutational burden. A) Distribution of DNA repair gene absences across the W/S clade and closest relatives. Presence/absence of *PHR1*, a gene encoding a DNA photolyase involved in UV damage repair is shown next to the phylogeny. The Outgroup 1 comprises species from the genera *Blastobotrys*, *Sugyamaella*, *Groenewaldozyma*, *Zygoascus*, *Trichomonascus*, and *Spencermartinsiella* and the Outgroup 2 comprises representative species from the genera *Magnusiomyces*, *Dipodascus*, *Geotrichum*, and *Saprochaete*. B) Analyses of substitution patterns among codon-based alignments of 143 single-copy orthogroups. Top row: Left) Substitution A|T bias. Y-axis is the number of substitutions in the A|T direction divided by the number of substitutions in the G|C direction. Thus, values greater than 1 indicate an A|T bias in substitutions. Right) ts/tv ratios reveal that the W/S clade is approaching a random expectation. Middle row) Mutational signatures associated with 8oxo-G, a common oxidative damage, reveal higher mutational signatures in the W/S clade. Bottom row: Single (T) and double (TT) mutations associated with UV damage. C) UV damage repair via Phr1 photolyase. UV damage generally results in pyrimidine (cytosine or thymine) dimers. The pruned species phylogeny was obtained from Opulente et al. (Opulente, et al. 2024). All pairwise statistical comparisons are shown with ns (no statistical significance), * (p-value < 0.05), ** (p-value < 0,001) or *** (p-value < 0.0001). All statistical analyses were performed using the Wilcoxon test with holm correction.

In the W/S clade, the genes absent among all species ranges from 32-55. In line with the slightly higher evolutionary rates in *Starmerella* compared to *Wickerhamiella* (Fig. S2A), we found that the average number of genes absent in *Starmerella* species (44) was slightly higher than in *Wickerhamiella* species (39) (Table S2). We also observed that a substantial portion of genes absent in the W/S clade are also absent in the W/S sister clade (Table S3). Specifically, 26 of the 37 genes absent in ≥ 50% of W/S-clade species were also absent in at least two of the three W/S- sister species (Table S3). Notwithstanding sampling issues associated with the small number of W/S-sister species, these results suggest that loss of DNA repair genes occurred both prior to and during the diversification of the W/S clade.

This prompted us to look for genes absent in the W/S clade and present in closest relatives, which would be strong candidates for being involved in accelerating evolutionary rates. We detected that *AHC1* was absent in all W/S clade species and present in the W/S-sister clade (Table S3). Ahc1 is part of the ADA complex involved in chromatin remodelling (Eberharter, et al. 1999), and loss-of-function mutations in this gene can cause elevated level of homologous recombination in *S. cerevisiae* (Wong, et al. 2013). *HNT3* and *ISY1* were also present in W/S sister and absent in most species of the W/S clade. Hnt3 is a DNA 5’-adenylate hydrolase involved in DNA damage response (Daley, et al. 2010), while Isy1 is involved in mRNA splicing (Villa and Guthrie 2005).

We noted that *CSM2* and *RFA3* were present in the W/S clade’s closest relatives and absent in most or all W/S-clade species. However, when we analyzed the sequences found in W/S sister species by a reciprocal BLASTp in NCBI non-redundant (nr) database, we concluded that these proteins were not orthologs of either *CSM2* or *RFA3*. *RFA3* is an essential gene in *S. cerevisiae* that encodes a DNA-binding subunit of replication protein A complex involved in DNA recombination (Brill and Stillman 1991). The putative Rfa3 sequences in W/S sister returned Hst4, an NAD^+^-dependent protein deacetylase involved in mitotic DNA replication and genomic stability (Pan, et al. 2006), as top hits, not Rfa3. As for *CSM2*, the top hits were Cdc40 proteins, which in *S. cerevisiae* encode a pre-mRNA splicing factor involved in cell cycle progression and RNA splicing (Dahan and Kupiec 2004). Although these proteins appear to belong to different gene families and perform distinct functions, they share sufficient homology to be detected by our searches. This overestimation of gene presences is likely the result of our use of a permissive e-value cutoff of 1e⁻³ in our sequence similarity searches, a decision guided by the need to prioritize confident inference of gene absences.

While our approach for identifying the absence of genes is conservative—employing an e-value cutoff of 1e⁻³ alongside tBLASTn searches against genome assemblies to mitigate the impact of annotation issues—the higher evolutionary rates observed in W/S-clade species may cause additional challenges in sequence homology detection. Specifically, some absences could stem from proteins whose sequences are too divergent to be recognized as homologs by sequence similarity search algorithms, a phenomenon previously documented in other highly divergent fungi (Mascarenhas Dos Santos, et al. 2022). To test whether this was an issue, we also employed a protein structural homology-based search that is not reliant on sequence similarity (Kaminski, et al. 2023). We chose five proteins absent in most W/S-clade species but present in their closest relatives (*AHC1*, *ISY1*, *HNT3*, *SLX1*, and *SLX4*). This approach confirmed the result of the sequence similarity searches, further suggesting the general absence of homologs to these DNA repair gene in the W/S clade (Table S5).

### Characterization of mutational patterns reveals a higher burden in the W/S clade

Our findings indicate that most absent genes in the W/S clade are also absent in the W/S-sister clade. However, while evidence of accelerated evolutionary rates was found in the W/S clade, no such evidence was found in its sister clade. Genomic fingerprints of base substitution patterns and indels, which provide insights into the mutational landscape, have previously been used to reveal signatures of mutational burden associated with DNA repair gene loss in *Hanseniaspora* FEL (Steenwyk, et al. 2019). Thus, we analyzed patterns of base substitutions, substitution directionality, indels, and signatures of endogenous and UV-induced damage across W/S clade species and their closest relatives (Fig. 3 and Fig. S6).

We found that the W/S clade exhibited a higher mutational burden than its closest relatives across several mutational signatures examined. For instance, the bias towards A|T substitutions was significantly stronger in the W/S clade than in its relatives (p-value < 0.001, Wilcoxon rank test), except when compared to the W/S-sister clade (Fig. 3B), a pattern consistent with the general A|T bias of mutations reported for several organisms (Hershberg and Petrov 2010; Lynch 2010; Liu and Zhang 2021), including the *Hanseniaspora* FEL (Steenwyk, et al. 2019). In line with this, the transition (ts)/transversion (tv) ratio was approximately 0.5–0.7 in the W/S clade (Fig. 3B). These values align with the estimated ts/tv ratios attributed to neutral mutations in *S. cerevisiae* (Lynch, et al. 2008; Zhu, et al. 2014). We also found significant mutational load associated with one of the most abundant endogenously damaged bases, 8-oxoguanine (Shibutani, et al. 1991), which causes the transversion mutation of G → T or C → A (De Bont and van Larebeke 2004).

Deletions were also significantly higher in the W/S clade compared to its closest relatives (Fig. S6), while no evidence for higher proportion of insertions was found (Fig. S6). We also did not find evidence for mutational burden associated with UV damage, which was evaluated by the number of C → T substitutions at CC sites (or G → A substitutions at GG sites), as well as the less frequent CC → TT (or GG → AA) double substitutions (Fig. 3B). In fact, we found that C → T substitutions in particular might be lower in the W/S clade when compared to closest relatives.

Interestingly, while we did not find evidence of accelerated evolutionary rates in the W/S-sister clade (Fig. 3A), this lineage showed slight bias towards A|T substitutions (Fig. 3B), suggesting a lower DNA repair efficiency. Consistent with this hypothesis, the ts/tv ratio in this clade was also notably lower than in other closely related clades (Fig. 3B). This finding suggests that, while loss of DNA repair genes might not have affected evolutionary rates in this lineage (Fig. 3A), it could have left mutational fingerprints across the genome. However, it is important to note that these considerations are based on only the three species currently described in the W/S sister clade.

### Horizontal acquisition of DNA repair genes may buffer gene losses

*PHR1* encodes a DNA photolyase and is the main gene responsible for directly reversing UV damage in *S. cerevisiae* (Liu, et al. 2011). Considering the absence of UV damage signatures in the W/S clade, we investigated the distribution of the *PHR1* gene within this lineage and closest relatives. Approximately 84% of species in the W/S clade possess a *PHR1* gene, whereas the gene is substantially less prevalent among their closest relatives (Fig. 3A).

The patchy distribution of *PHR1* spurred us to investigate its evolutionary history in the W/S clade. For that, we performed a BLASTp search of all putative Phr1 sequences from the W/S clade against the NCBI refseq database. For some W/S-clade species, the top hits were from filamentous fungi (subphylum Pezizomycotina); for some *Wickerhamiella* species, the top hits were bacterial proteins. This suggests that HGT event(s) might have contributed to the patchy distribution of Phr1 in the W/S clade. To formally test this, we constructed a phylogenetic tree with all putative orthologs of Phr1 from Saccharomycotina (using the 1,154-proteome dataset) and the top hits obtained from refseq for two distinct W/S clade Phr1 proteins (bacterial and fungal). The phylogeny, represented in Figure 4A, shows three major clades: one of bacterial sequences, one of Saccharomycotina sequences, and a third of Pezizomycotina sequences; the Saccharomycotina and Pezizomycotina sequences were sister clades, as expected from the species phylogeny (Li, et al. 2021). There were two exceptions to this pattern. First, some W/S-clade Phr1 proteins cluster within the bacterial clade (within the order Sphingomonadales), and the rest of the W/S-clade Phr1 proteins clustered within the Pezizomycotina clade. The second exception were sequences of *Blastobotrys* species (Outgroup 1), which clustered within the Pezizomycotina clade (together with W/S Phr1 sequences) (Fig. 4A). Constrained topology analyses rejected an alternative topology where bacterial-like W/S sequences clustered with other Saccharomycotina species (p-value = 0.001, Approximately Unbiased [AU] test). This result supports the hypothesis that an HGT event occurred, possibly in the most recent common ancestor (MRCA) of a *Wickerhamiella* subclade (Fig. 4B). As for the Pezizomycotina-like Phr1 sequences from the W/S clade, constrained topology tests also rejected the alternative hypothesis that clustered these proteins with other Saccharomycotina (p-value = 0, AU test). Nevertheless, most Pezizomycotina-like W/S Phr1 sequences formed a monophyletic group, which is sister to the Pezizomycotina. Therefore, we cannot exclude the alternative hypothesis of differential retention of an ancestral paralog.

**Figure 4.**
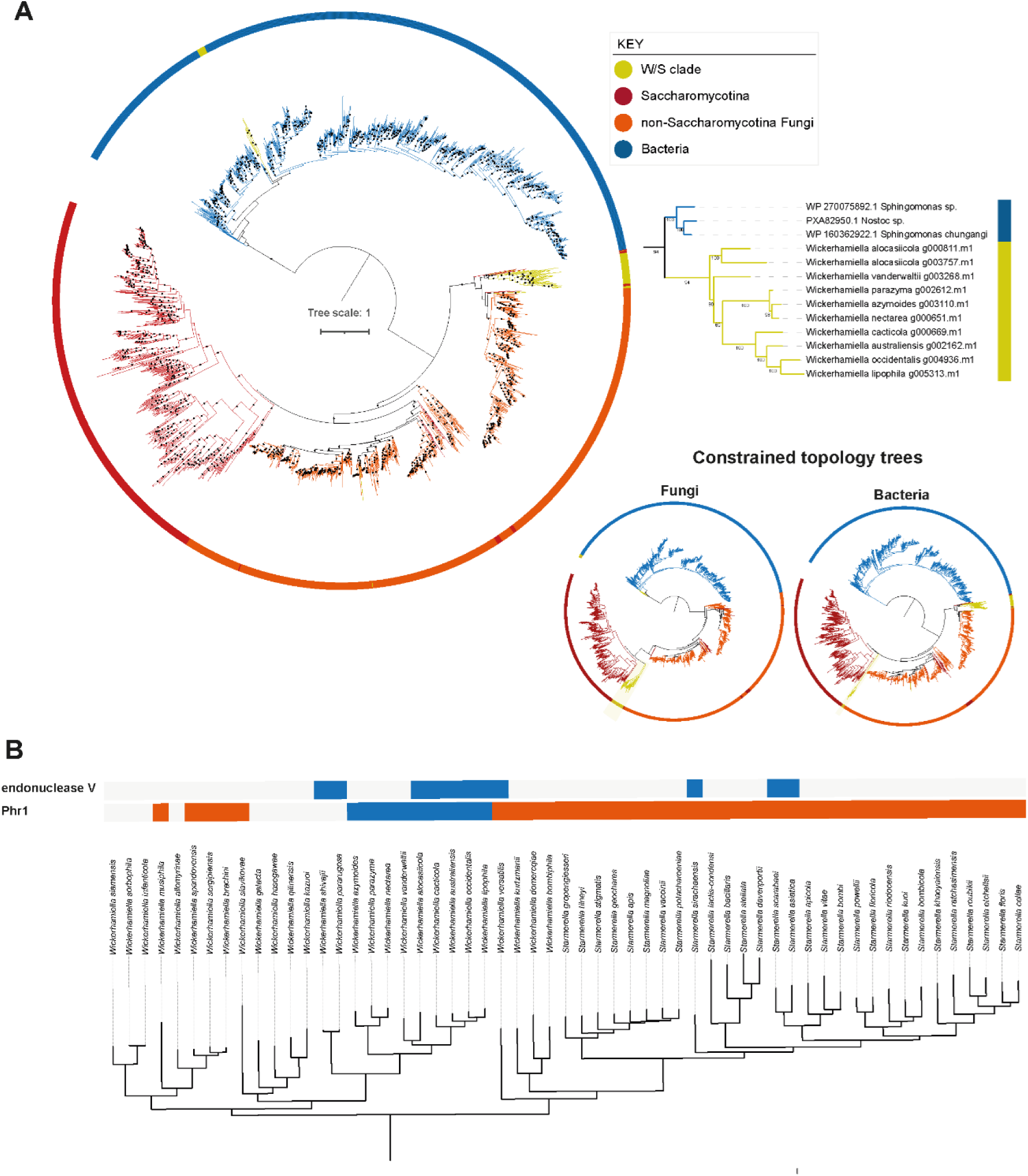
Several W/S-clade species acquired bacterial DNA repair genes through horizontal gene transfer (HGT). A) Phylogenetic tree of sequences with the highest sequence similarity to Phr1 from *Wickerhamiella australiensis* and *Starmerella bombicola*. (right panel) Pruned Phr1 phylogeny depicting the HGT event from bacteria (Sphingomonadales) to a W/S subclade. On the bottom, the resulting trees from the constrained topology analyses are shown: Fungi – constrained topology considering monophyly of the W/S Pezizomycotina-like sequences and other Saccharomycotina (excluding the Saccharomycotina sequences that cluster with other Fungi) and Bacteria-constrained topology considering monophyly of bacteria-like W/S sequences and Saccharomycotina sequences (excluding those clustering with other fungi). B) Distribution of putative HGT-derived (blue: bacterial, orange: fungal) *PHR1* and endonuclease V genes across the W/S clade.

We next tested whether these putative xenologs are functional by performing UV irradiation assays (Milo, et al. 2019). We found that, in W/S-clade species harbouring bacterial and fungal Phr1 versions, cells pre-exposed to UV-A light (photoreactivation, low damage to DNA) were more resistant to UV-C (which causes lesions in DNA) than cells that were not subject to photoreactivation (Fig. S7). This is consistent with the UV-A dependent activation of Phr1 (Sancar 1990; Milo, et al. 2019).

It is well established that the W/S clade has acquired numerous foreign genes from both bacteria and filamentous fungi (Gonçalves, et al. 2016; Gonçalves, et al. 2018; Shen, et al. 2018; Gonçalves and Gonçalves 2019; Kominek, et al. 2019; Gonçalves, et al. 2020; Pontes, et al. 2024). To explore if other DNA repair-related genes might have also been acquired horizontally by W/S-clade species, we retrieved the DNA repair genes from *Escherichia coli* and *Bacillus subtilis* (total of 263 sequences) as of February 2023 and performed a BLASTp against the W/S proteomes. We next constructed phylogenetic trees for the proteins that showed significant similarity with bacterial proteins based on a second BLAST search against the NCBI nr database. From this analysis, we detected seven DNA repair-related proteins whose top BLASTp hits were bacterial sequences (Table S4), suggesting that they are of putative bacterial origin. One of these proteins is a homolog of *MGT1*, which appears to have also been involved in HGT events in *Hanseniaspora* FEL and *C. sorboxylosa* (Pichiales subclade) (Fig. S5). Phylogenetic reconstruction provided support for the bacterial origin of an endonuclease V, which was likely independently acquired by W/S-clade species (Fig. 4B), possibly once in the MRCA of *Wickerhamiella* and once in the MRCA of *Starmerella* (Fig. S8A). Although top BLAST hits for *Wickerhamiella* species were endonuclease V sequences from the bacterial CFB group, the *Starmerella* top hits belonged to the gamma proteobacteria. This topology suggests at least two independent acquisitions, but we did not find statistical support for this hypothesis (p-value = 0.069, AU test). Endonuclease V is involved in the repair of deaminated DNA bases, which are commonly caused by endogenous and environmental agents (Cao 2013) and is absent in the rest of the Saccharomycotina. However, loss of endonuclease V in *Schizosaccharomyces pombe* (subphylum Taphrinomycotina) induces a strong mutator phenotype (Dalhus, et al. 2009). Another protein for which we found phylogenetic support of HGT was UvrA (Fig. S8B), which is partly involved in nucleotide excision repair and might also be involved in UV damage repair in bacteria (Agostini, et al. 1996; Crowley, et al. 2006).

## Discussion

Rapid evolution associated with the loss of DNA repair genes is usually observed in microevolutionary contexts, such as during the evolution of human tumours or clinical isolates of microbial pathogens (LeClerc, et al. 1996; Fukui 2010; Healey, et al. 2016; Billmyre, et al. 2017; Campbell, et al. 2017; Gambhir, et al. 2022). A few examples demonstrate how entire fungal lineages can also undergo rapid evolution over macroevolutionary timescales, likely resulting from the loss of DNA repair genes (Steenwyk, et al. 2019; Phillips, et al. 2021). Here, we demonstrate that the link between rapid evolution and impairment of DNA repair holds for the three most rapidly evolving lineages of the fungal subphylum Saccharomycotina. One of these lineages, belonging to the genus *Hanseniaspora* (*Hanseniaspora* FEL), was previously identified as having lost numerous genes associated with the cell cycle and DNA repair (Steenwyk, et al. 2019), demonstrating the efficacy of our approach, while the other two lineages, the *Wickerhamiella*/*Starmerella* (W/S) clade and a subclade within the Pichiales comprising *Pichia, Saturnispora*, and *Martiniozyma* species are reported here for the first time.

The number of absent genes in these three FELs is significantly higher than in the rest of the subphylum; however, we found that few yeasts have a “complete” DNA repair gene repertoire. This phenomenon can be partially explained by the fact that most of the genes inspected are part of the *S. cerevisiae* gene repertoire and supports the notion that the DNA repair gene repertoire varies across yeast species (Milo, et al. 2019). It also indicates that many of these genes are not essential and may be partially redundant, which is a critical feature in safeguarding the genome against DNA damage (Darzynkiewicz 2011; Gartner and Engebrecht 2021). Even genes described as essential in *S. cerevisiae* and close relatives, such as the transcription regulator Abf1 (Rhode, et al. 1992; Hernández-Hernández, et al. 2021) might not be essential in the distinct genetic (and environmental) backgrounds of other species in the Saccharomycotina subphylum. For instance, insects are known to lack several “key” DNA repair genes, suggesting that they may have found alternative strategies to deal with DNA-damaging agents (Wyder, et al. 2007; Sekelsky 2017).

Although we found that the lineages with the most reduced DNA repair gene repertoires were also the fastest evolving, the correlation between the size of the DNA repair gene repertoire and rates of evolution across the subphylum was weak. One potential explanation for the weak correlation is that high mutation rates can be caused by only a few genes rather than being inversely proportional to the size of the DNA repair gene repertoire. In line with this hypothesis, we sought to identify genes potentially involved in increasing evolutionary rates by analysing the common absences across the three FELs. We did not find genes exclusively absent in FELs, but several genes (e.g., *MAG1*, *MGT1*) were absent in most of the faster-evolving species, suggesting that they might be or have been involved in acceleration of mutation rates in these genetic backgrounds.

In line with the weak correlation between DNA gene repertoire size and evolutionary rates, we found that the most significant loss of DNA repair-related genes might have occurred before the diversification of the W/S clade since its sister clade already exhibits reduced DNA repair gene repertoires. We observed that most genes absent in the W/S clade are also absent in species belonging to its sister clade, but we did not find evidence of accelerated evolutionary rates in the latter. However, the W/S-sister clade does display mild signatures of mutational burden, such as A|T bias. This discrepancy could either suggest that the additional genes losses in the W/S clade (e.g., *AHC1*, *ISY1*) may be involved in the acceleration of the evolutionary rates or that the W/S- sister clade may have compensated the loss of certain DNA repair genes through alternative mechanisms.

Interestingly, we found possible mechanisms of compensation for the loss of certain DNA repair genes in the W/S clade, such as through the acquisition of foreign genes associated with UV damage repair. We found that most W/S-clade species encode Phr1 proteins, while most of their closest relatives lack the gene. Some of the W/S-clade Phr1 homologs were likely acquired from bacteria in a single HGT event, while the others are more closely related with Phr1 sequences from filamentous fungi. Whether the fungal-like sequences are the result of additional HGT event(s) from the Pezizomycotina or are the result of a differential retention of an ancient paralog remains unclear.

In line with the presence of bacterial or fungal Phr1 sequences in almost all W/S-clade species, we found no evident genomic signatures of UV-induced damage and that Phr1-containing species are resistant to UV under photoreactivation conditions, which suggests that these proteins are functional. Additionally, we found that some species also acquired a bacterial endonuclease V, a broad specificity enzyme involved in the repair of deaminated bases that can arise from multiple types of endogenous and exogenous aggressions (Cao 2013). Although species from the W/S clade seem to have found alternative pathways to deal with DNA damage, it is still unclear whether a deceleration of the evolutionary rates have occurred, as observed for *Hanseniaspora* (Steenwyk, et al. 2019). Mutation accumulation experiments will be essential to ascertain whether these species are still evolving faster or have slowed down their mutation rates.

It is important to note that our conservative approach to considering gene absences may generate false positives (false gene presences) by detecting the presence of conserved motifs in distant homologs as shown for *CSM2* or *RFA3*. Conversely, reduced DNA repair gene repertoire was previously associated with remarkably high levels of sequence divergence in a lineage of intracellular parasites (Microsporidia) (Corradi 2015). However, using a recently developed protein language model for distant homolog detection, DNA repair gene loss in Microsporidia was found to have been less extensive than previously thought (Mascarenhas Dos Santos, et al. 2022). Although we validated some of the gene absences with both tBLASTn searches against the genomes to avoid annotation issues and (in a few cases) with non-similarity-based methods for detecting distant homology, we cannot rule out that some genes are fast-evolving and are therefore difficult to identify. However, at least in the W/S clade, loss of DNA repair genes seems to have been part of a broader gene loss event (Fig. S9). While *Blastobotrys* and *Sugiyamaella* yeast genomes range between 11–25 Mb in size, W/S-clade genomes are around 9-11 Mb. In line with the observation that the W/S sister clade also lost a significant portion of their DNA repair gene repertoire (Fig. 3A), we observed a slight decrease in both genome size and number of CDS in this lineage, suggesting that extensive ancient gene losses might have occurred in the MRCA of the W/S and W/S-sister clades, with subsequent gene loss events occurring after the diversification of the W/S clade, including genes involved in DNA repair. This suggests that some of the HGT events of DNA repair genes we uncovered, and others documented in the literature (Gonçalves, et al. 2018; Gonçalves and Gonçalves 2019; Gonçalves, et al. 2022; Pontes, et al. 2024), might have been compensatory for these ancient gene losses.

The W/S clade is in fact known for its very high numbers of horizontally acquired genes (Gonçalves, et al. 2018; Gonçalves and Gonçalves 2019; Kominek, et al. 2019; Gonçalves, et al. 2020; Pontes, et al. 2024), exhibiting the highest number of bacterial genes across the subphylum Saccharomycotina (Shen, et al. 2018). Defects in certain DNA repair pathways, such as the MMR system, have been correlated with higher rates of HGT events in bacteria because these mutator strains can recombine more frequently with divergent DNA (Rayssiguier, et al. 1989; Thomas and Nielsen 2005). While we did not find specific MMR losses in the W/S clade, we can speculate whether the periods of high mutation rates might have facilitated the integration of HGT-derived genes into W/S yeast genomes.

In conclusion, while the loss of DNA repair genes may be generally detrimental due to the accumulation of deleterious mutations, our results show that entire lineages can maintain elevated mutation rates for prolonged periods of time. In the well described short-lived hypermutator populations (Billmyre, et al. 2017; Rhodes, et al. 2017; Gambhir, et al. 2022; Callens, et al. 2023; Hall, et al. 2025) high mutation rates are associated with nonsense mutations in DNA repair genes that can be compensated by the emergence of anti-mutator alleles. Over macroevolutionary timescales, genetic constraints stemming from wholesale gene loss of DNA repair genes may contribute to the maintenance of high evolutionary rates. These results also highlight that these genetic constraints can be overcome by mechanisms such as HGT of DNA repair genes.

## Materials and methods

### Gene presence and absence analysis and determination of evolutionary rates

To assess the presence and absence of DNA repair genes across the Saccharomycotina subphylum, we first retrieved all DNA repair proteins belonging to Saccharomycotina yeasts from UniprotKB (keywords: dna+repair+saccharomycotina-filtered-reviewed) as of February 2023. Although we searched for all Saccharomycotina DNA repair related proteins at UniprotKB, most of the retrieved genes belong to *Saccharomyces cerevisiae* (Table S1). These 415 proteins are associated with distinct DNA repair functions from DNA damage response, cell cycle, chromatin or telomere organization (Table S1). For each protein, we built HMMs profiles (Eddy 1998) using an alignment with the top 100 hits retrieved by BLASTp from the NCBI non-redundant (nr) database. Next, we ran Orthofisher (Steenwyk and Rokas 2021) with default parameters (e-value cutoff < e^-3^) and with higher levels of stringency (e-value cutoff of 1e^-20^ and 1e^-50^) using the previously constructed HMM profiles against 1,154 proteomes corresponding to 1,090 Saccharomycotina species (Opulente, et al. 2024).

Absences were confirmed for the common absent genes in the three FELs (Fig. 2) and for the list of genes absent in the W/S clade (Table S3) by tBLASTn searches against the respective assemblies using the list of 415 DNA repair related proteins. Genes were considered absent whenever the e-value for the best hit was > 0.1. For the hits with e-values < 0.1, identity of the gene was confirmed BLASTp against the NCBI nr database using *Saccharomyces cerevisiae* as reference.

Evolutionary rates were determined using tip-to-root distances as proxy. For that we used the Saccharomycotina phylogeny constructed in Opulente et. al., 2024 (Opulente, et al. 2024) and determined the tip-to-root distances using the *distRoot* function (method=patristic) included in the *adephylo* package for R (Jombart, et al. 2010).

### Large Language Model (LLM) remote homolog detection

To search for more diverged homologs without relying on sequence similarity we used pLM-BLAST, a large language model (LLM) base approach optimized for remote homolog detection (Kaminski, et al. 2023). To prepare to run pLM-BLAST, we first used the pLM-BLAST tool embeddings.py to make protein sequence embeddings of each of the genes, in all the proteomes of the W/S clade (307,448 sequences in total). We then used embeddings.py to make embeddings for representative proteins in the sister clade (*Blastobotrys* group) (45,011 sequences, clustered by mmseq2 (Steinegger and Söding 2017) to represent the 450,863 proteins in the clade). Embeddings were then generated for each query sequence (*AHC1* from *Candida incommunis*, *HNT3* from *Sugiyamaella lignohabitans*, *ISY1* from *Candida incommunis*, *SLX1* from *Sugiyamaella lignohabitans*, and *SLX4* from *Sugiyamaella lignohabitans*). Next, each query embedding was searched using the default pLM-BLAST parameters (alignment cutoff =0.3, cosine cutoff=90, sigma=2) against all proteins in the W/S and sister clade, using on 4x NVIDIA RTX A6000 GPUs. Finally, pLM-BLAST hits were sorted by their cosine similarity score, and any queries having scores above 0.5 were considered as possible homologs (Table S5). For hits with scores higher than 0.5, a subsequent reciprocal BLASTp against NCBI nr database (using *Saccharomyces cerevisiae* as reference) was performed. These results were mostly negative, with most hits being fragmentary and likely corresponding to individual domains rather than whole genes. Overall, these remote homolog detection results recapitulate the results of the sequence similarity searches.

### Mutational signatures

To identify base substitutions and indels (insertions and deletions) in the W/S clade, we followed previously published methodologies (Steenwyk, et al. 2019). First, we select the orthologues to be analysed by running OrthoFinder v.2.3.8 (Emms and Kelly 2019) using an inflation parameter of 1.5 and DIAMOND v2.0.13.151 (Buchfink, et al. 2015) for protein alignments, on a dataset containing all W/S-clade proteomes (excluding redundancy, i.e., strains from the same species) as well as proteomes from closest relatives (W/S sister, Outgroups 1 and 2) and outgroups (Table S6). A total of 143 core single-copy orthogroups were obtained and subsequently aligned with MAFFT v7.402 using an iterative refinement method (--localpair) (Katoh and Standley 2014; Vialle, et al. 2018). Next, codon-aware alignments for each amino acid alignment were generated with PAL2NAL v.14 (Suyama, et al. 2006).

Substitution patterns (a proxy for mutational patterns) were examined from the resulting multiple sequence alignments. To do so, substitutions, insertions, and deletions were examined at sites otherwise conserved in the outgroup of closely related taxa (for example, the W/S sister clade is the outgroup to the W/S clade). For substitutions, the nucleotide character for a focal species was compared to the conserved nucleotide in the outgroup taxa at the same position. If the focal species had a nucleotide character that differed from the outgroup taxa, a substitution was determined to have occurred. While doing so, we kept track of the nucleotide character for the focal species and the outgroup taxa as well as the codon position, enabling inference of the directionality of the substitution and positional information. To ensure the number of mutations were comparable for each group, the raw number of substitutions were corrected by the number of conserved sites in the outgroup taxa. The same correction was made for substitutions at the various codon positions. For the AT-bias analysis, a correction was made to account for variation in the conserved number of GC or AT sites. Lastly, a correction was made to normalize for the number of single-copy orthologous genes that could be examined per focal lineage and closely related lineage. To identify insertions and deletions, a sliding window approach with a step size of one nucleotide was used to scan the multiple sequence alignments for positions that had nucleotides in the focal species and gaps in the outgroup taxa (insertions) or vice versa (deletions).

Statistical significance of the differences between the several groups (W/S clade, W/S sister, Outgroup 1 and Outgroup 2) were assessed by pairwise Wilcoxon rank test after testing for normality.

### Phylogenetic analyses of *PHR1*

Putative Phr1 sequences from W/S-clade species and other Saccharomycotina were retrieved from the Orthofisher run and used in BLASTp searches against the NCBI nr database. Two distinct major lineages were identified as top hits for different W/S-clade species: Pezizomycotina (filamentous fungi) and Bacteria. Putative Phr1 sequences from *Wickerhamiella australiensis* (top hits Bacteria) and *Starmerella bombicola* (top hits Pezizomycotina) were used in a BLASTp search against the NCBI refseq database (O’Leary, et al. 2015). The top 250 hits from each blast search were retrieved. Redundancy was removed with CD-HIT (keeping sequences with less than 98% identity) (Li and Godzik 2006). The resulting 2,007 sequences and subsequently aligned using MAFFT v7.402 using an iterative refinement method (--localpair) and poorly aligned regions were removed with trimAL (Capella-Gutierrez, et al. 2009) using the “gappyout” option. A Phr1 phylogeny was subsequently constructed with IQ-TREE v2.0.6 (Nguyen, et al. 2015) using an automated method for model selection and 1,000 ultrafast bootstraps (Hoang, et al. 2018). The tree with the highest likelihood score was subsequently chosen from a total of five runs (--runs 5). The hypothesis of the horizontal acquisition of Phr1 from both bacteria and filamentous fungi was assessed through topology tests performed in IQ-TREE v2.0.6 (Nguyen, et al. 2015). For that, two constrained topologies were constructed. In the first constrained topology to test HGT from bacteria, W/S-clade sequences clustering with bacteria were considered to be monophyletic with other Saccharomycotina (except for Saccharomycotina sequences that clustered with other fungi). To test for HGT from filamentous fungi, a second constrained topology was constructed by considering “fungal” W/S-clade sequences as monophyletic with other Saccharomycotina (except for Saccharomycotina sequences that clustered with other fungi). Constrained trees were constructed in IQ-TREE v2.0.6 (using the “-g” option). The likelihoods of the constrained and unconstrained tree topologies were subsequently compared in IQ-TREE using the AU test (using “-z” option).

### UV sensitivity assays

Single colonies from three putative Phr1 positive W/S-clade yeast, *Wickerhamiella vanderwaltii*, *Wickerhamiella cacticola*, and *Starmerella sirachaensis* were resuspended in 200 μL water in a 96 well plate together with *Saccharomyces cerevisiae* (Phr1 positive) and *Brettanomyces bruxellensis* (Phr1 negative) (Milo, et al. 2019). The cultures were 10-fold serially diluted and spotted onto YPD [1% (w/v) yeast extract, 2% (w/v) bacto peptone and 2 % (w/v) glucose] agar plates. The plates were irradiated by 100 or 200 J/m^2^ using UV-C with or without two hours of photo-reactivation using an UV-A lamp as previously described (Milo, et al. 2019; Milo, et al. 2024).

### Search for additional bacteria-derived DNA repair genes

To investigate whether other DNA repair genes from bacteria were horizontally acquired by W/S-clade species, DNA repair genes from *Escherichia coli* K12 reference strain and *Bacillus subtilis* reference strain were retrieved from UniprotKB (as of February 2023). A local BLASTp search against all W/S-clade proteomes was performed (e-value cutoff 1e^-3^). The top blast hit for each gene was subsequently retrieved and analysed through a BLASTp search against the NCBI nr database and whenever the best hit corresponded to a bacterial gene, a phylogenetic tree was reconstructed to confirm the bacterial origin of the gene. Phylogenies were constructed as by retrieving the closest related sequences obtained by BLASTp searches against NCBI nr or UniprotKB databases. Sequences were aligned using MAFFT v7.402 using an iterative refinement method (--localpair) and trees were constructed with IQ-TREE v2.0.6 using an automated method for model selection and 1,000 ultrafast bootstraps.

## Acknowledgements

We thank members of the Rokas Lab, Hittinger Lab, Yeast Genomics lab and members of the Y1000+ Project (http://y1000plus.org) for helpful discussions. This work was performed using resources contained within the Advanced Computing Center for Research and Education at Vanderbilt University in Nashville, TN.

## Funding

National Science Foundation Grant DEB-2110403 (CTH) National Science Foundation Grant DEB-2110404 (AR)

DOE Great Lakes Bioenergy Research Center, funded by BER Office of Science Grant DE-SC0018409 (CTH)

USDA National Institute of Food and Agriculture Hatch Project 1020204 (CTH) USDA National Institute of Food and Agriculture Hatch Project 7005101 (CTH)

H. I. Romnes Faculty Fellow, supported by the Office of the Vice Chancellor for Research and

Graduate Education with funding from the Wisconsin Alumni Research Foundation (CTH)

National Institutes of Health/National Institute of Allergy and Infectious Diseases Grant R01 AI153356 (AR)

Burroughs Wellcome Fund (AR)

Research supported by the National Key R&D Program of China Grant 2022YFD1401600 (XXS)

National Science Foundation for Distinguished Young Scholars of Zhejiang Province Grant LR23C140001 (XXS)

Fundamental Research Funds for the Central Universities Grant 226-2023-00021 (XXS) National Institutes of Health Grant T32 HG002760-16 (JFW)

National Science Foundation Grant Postdoctoral Research Fellowship in Biology 1907278 (JFW)

Fundação para a Ciência e a Tecnologia Grant PTDC/BIA-EVL/0604/2021 (CG) Fundação para a Ciência e a Tecnologia Grant UIDP/04378/2020 (CG) Fundação para a Ciência e a Tecnologia Grant UIDB/04378/2020 (CG) Fundação para a Ciência e a Tecnologia Grant LA/P/0140/2020 (CG)

JLS is a Howard Hughes Medical Institute Awardee of the Life Sciences Research Foundation.

