## Supplementary Figures for "Stable hypermutators revealed by the genomic landscape of DNA repair genes among yeast species"

### Supplemental Figures

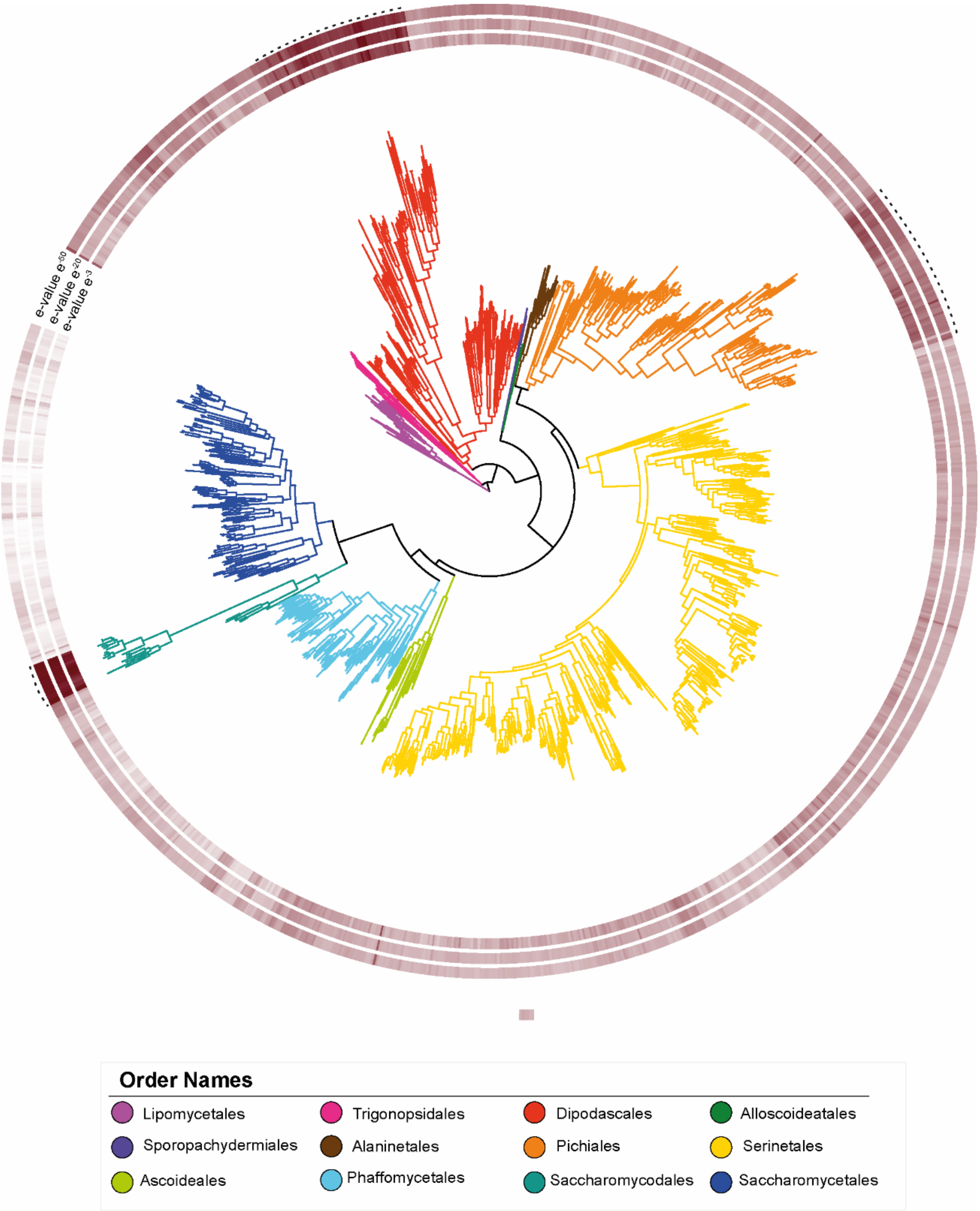


**Figure S1. Number of absent DNA repair genes across the Saccharomycotina using distinct e-value thresholds.** Distribution of the number of absent DNA repair genes (from a total of 415) across the Saccharomycotina species phylogeny (Opulente, et al. 2024). Distinct e-value thresholds were used representing distinct degrees of stringency for considering gene presence ($e^{-3}$, $e^{-20}$, $e^{-50}$). The distribution highlights that the three faster-evolving lineages (W/S clade, Pichiales subclade, and *Hanseniaspora* FEL) show a significant decrease in their DNA repair gene repertoire show the independently of the e-value threshold used.

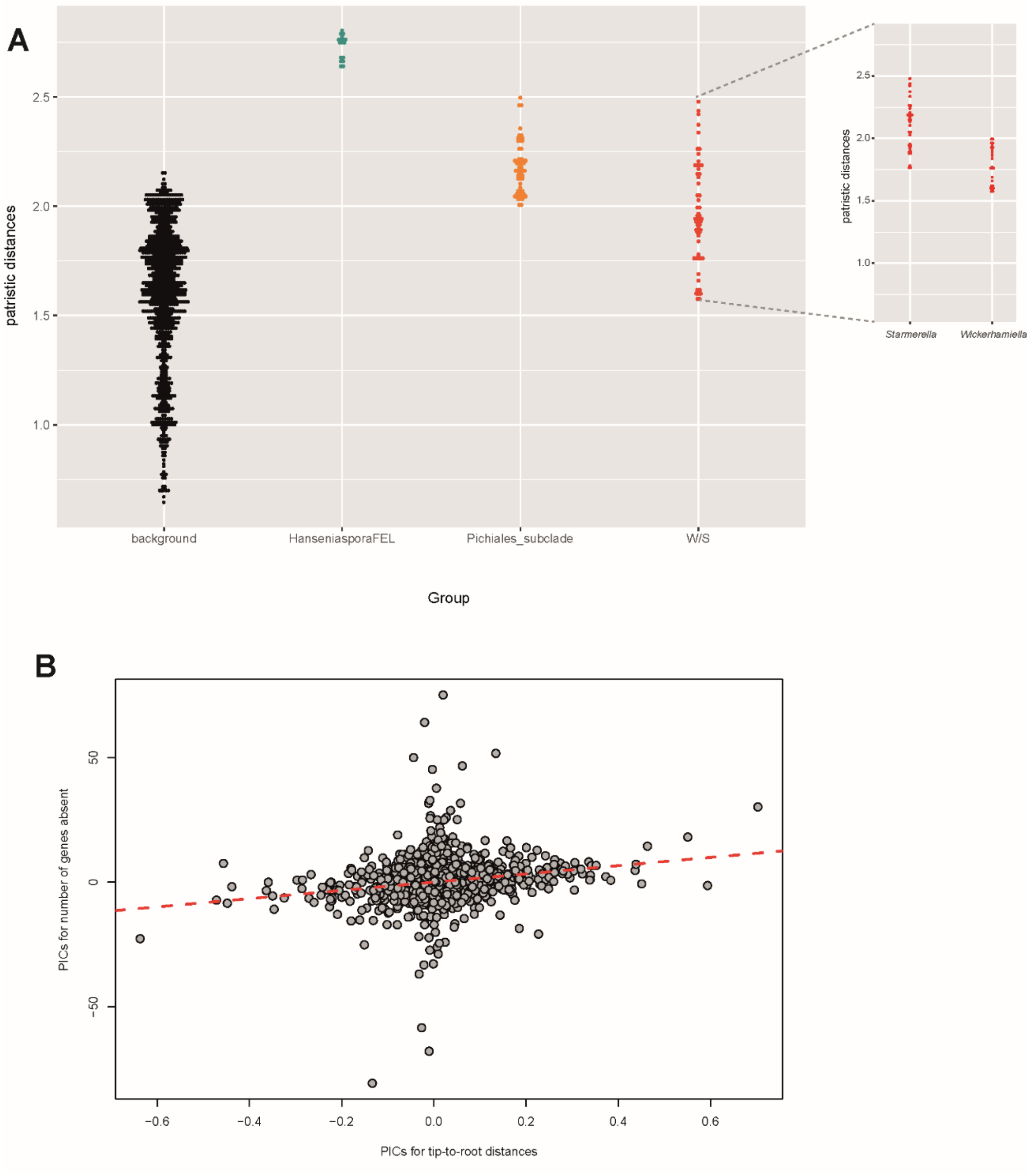


**Figure S2. Correlation between DNA repair repertoire and evolutionary rates. A)** Evolutionary distances in the three faster evolving lineages, highlighting the slight differences between *Starmerella* and *Wickerhamiella* genera in the W/S clade. B) Phylogenetically corrected (PIC) correlation between tip-to-root distances and number of absent DNA repair genes.

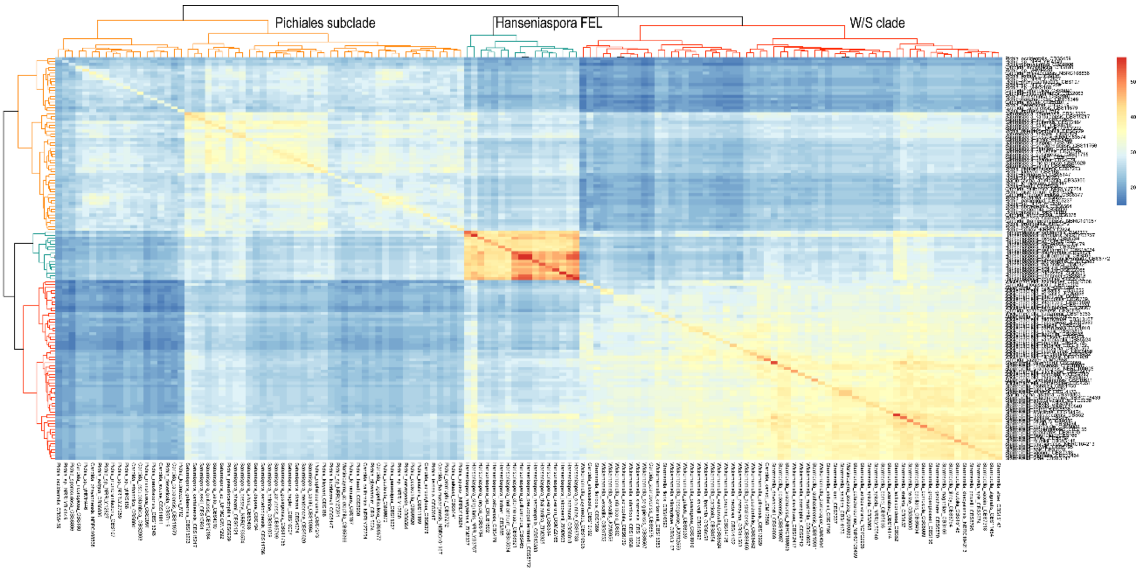


**Figure S3. Heatmap comparing the repertoire of DNA repair genes absent across the three faster evolving lineages.** The heatmap highlights the partial overlap of gene absences between the three lineages.

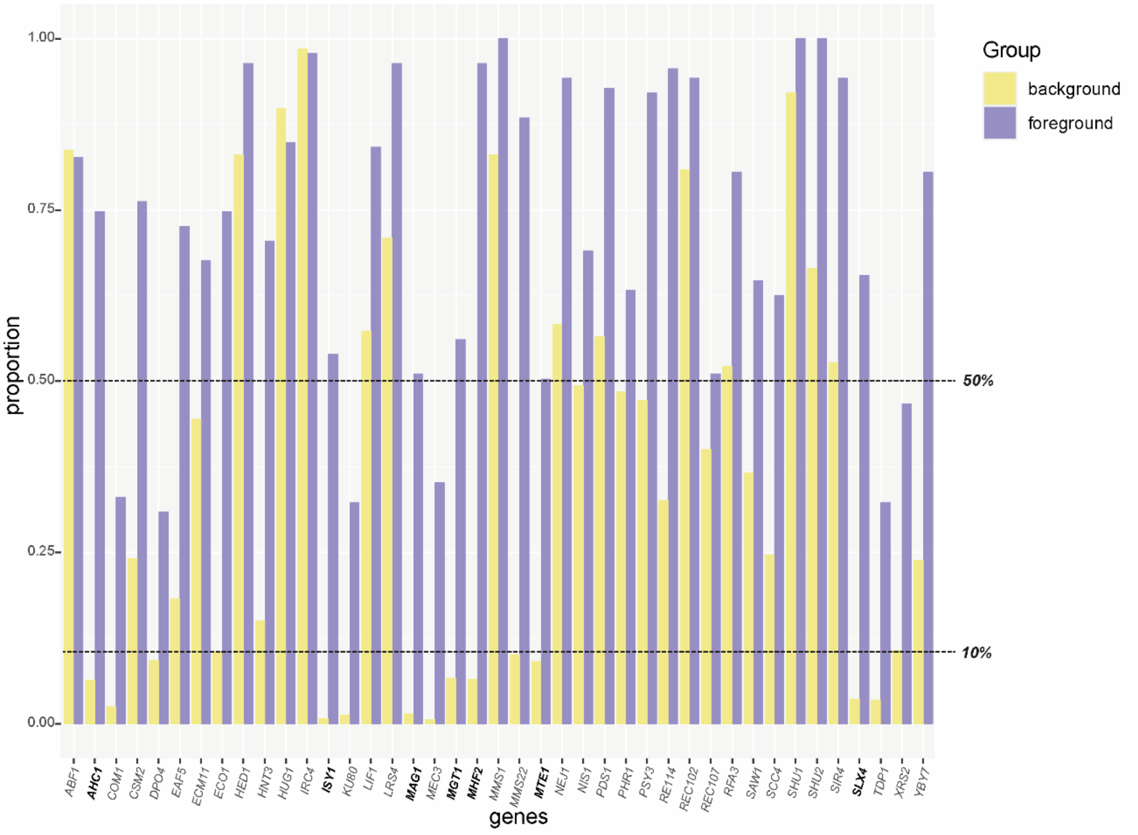


**Figure S4. Proportion of foreground (W/S clade, *Hanseniaspora* FEL, and Pichiales subclade) and background (all the other) species missing DNA repair genes.** Only genes that were absent in more than 30% of the species belonging to each faster evolving lineage were considered for this analysis. The proportions were then determined taking into account the three FELs all together (N=139 proteomes).

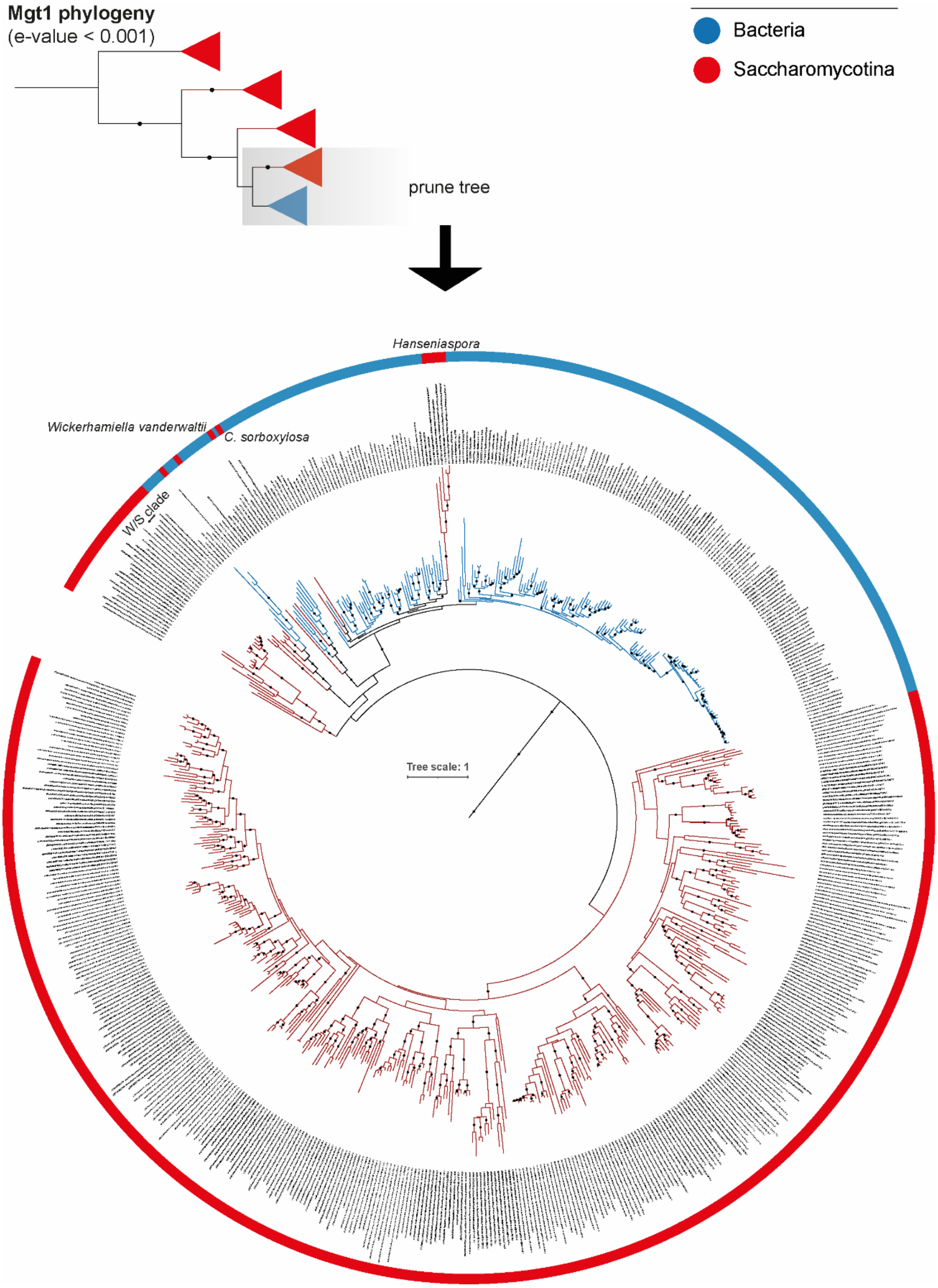


**Figure S5. Mgt1 phylogeny depicting the clustering of *Hanseniaspora* and *Candida sorboxylosa* sequences with those of bacteria.** All possible Mgt1 homologs were retrieved from orthofisher (p-value < 0.001). Closest related sequences (bacterial) to *C. sorboxylosa* and *Hanseniaspora* FEL were also retrieved and used to construct a phylogeny. Topology analysis (AU test) was assessed separately for *Hanseniaspora* and *C. sorboxylosa* considering their monophyly with other Saccharomycotina Mgt1. This analysis revealed that the constrained trees are not statistically significant less likely than the unconstrained tree (p-value > 0.5, AU test).

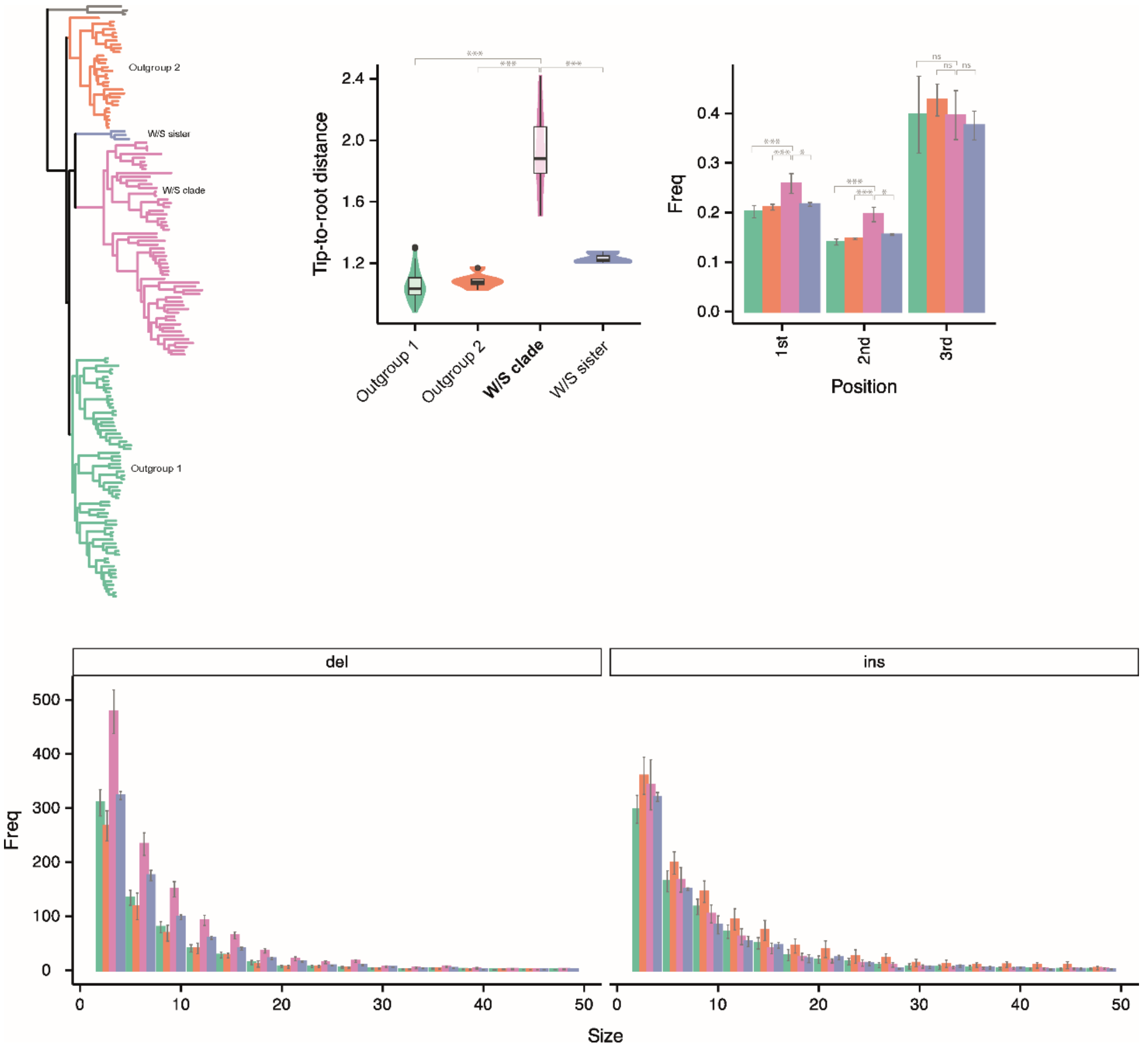


**Figure S6. Signatures of mutational burden across W/S clade and closest relatives.** Top row: Left) Tip to root distances; Right) Substitution frequency at different codon positions. Bottom left) Deletions of different codon sizes. Bottom right) Insertions of different codon sizes. Statistical significance was assessed using the Wilcoxon rank test after testing for normality.

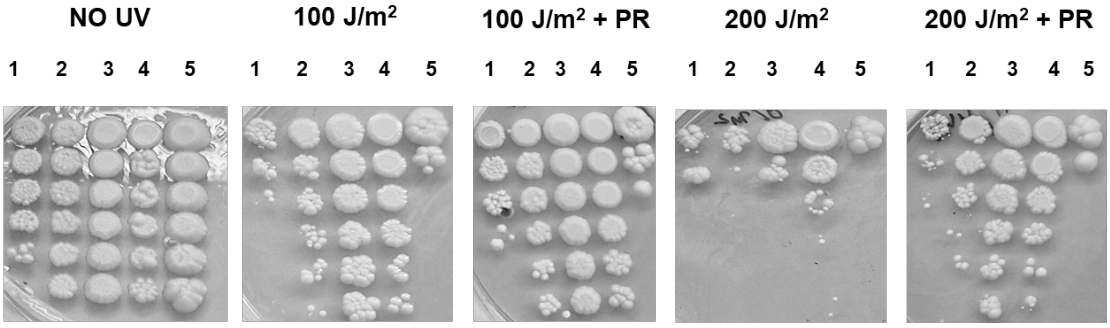


**Figure S7. UV sensitivity assays on plate (serial dilutions from top to bottom).** Yeast strains were spotted on YPD plates and irradiated with the indicated dose ($J/m^2$) with or without 2 hours of photoreactivation (PR). 1. *Saccharomyces cerevisiae* (control Phr1+); 2. *Wickerhamiella vanderwaltii* (encoding a bacterial Phr1); 3. *Wickerhamiella cacticola* (encoding a bacterial Phr1); 4. *Starmerella sirachaensis* (encoding a fungal Phr1); 5. *Brettanomyces bruxellensis* (control Phr1-).

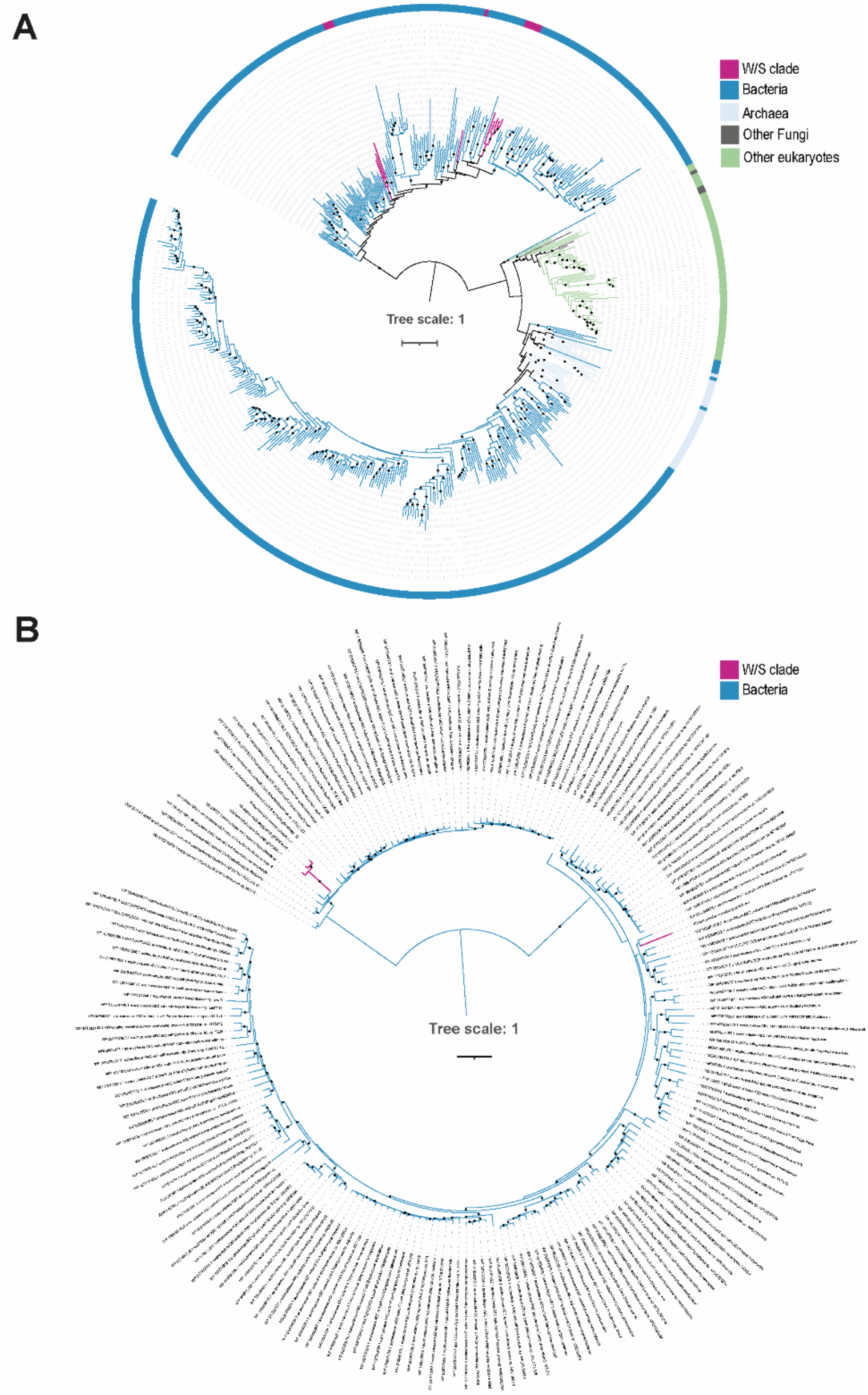


**Figure S8. Phylogenies of bacterial DNA repair related genes in the W/S clade.** A) Phylogeny of endonuclease V depicting the W/S clade grouping with distinct bacterial clades. B) *uvrA* phylogeny constructed using to top 500 BLASTp hits against NCBI nr database using *Wickerhamiella brachini* g002827.m1 protein as query. Redundancy was removed using CD-HIT (similarity> 98%), sequences were aligned using MAFFT and phylogeny was constructed in IQ-TREE.

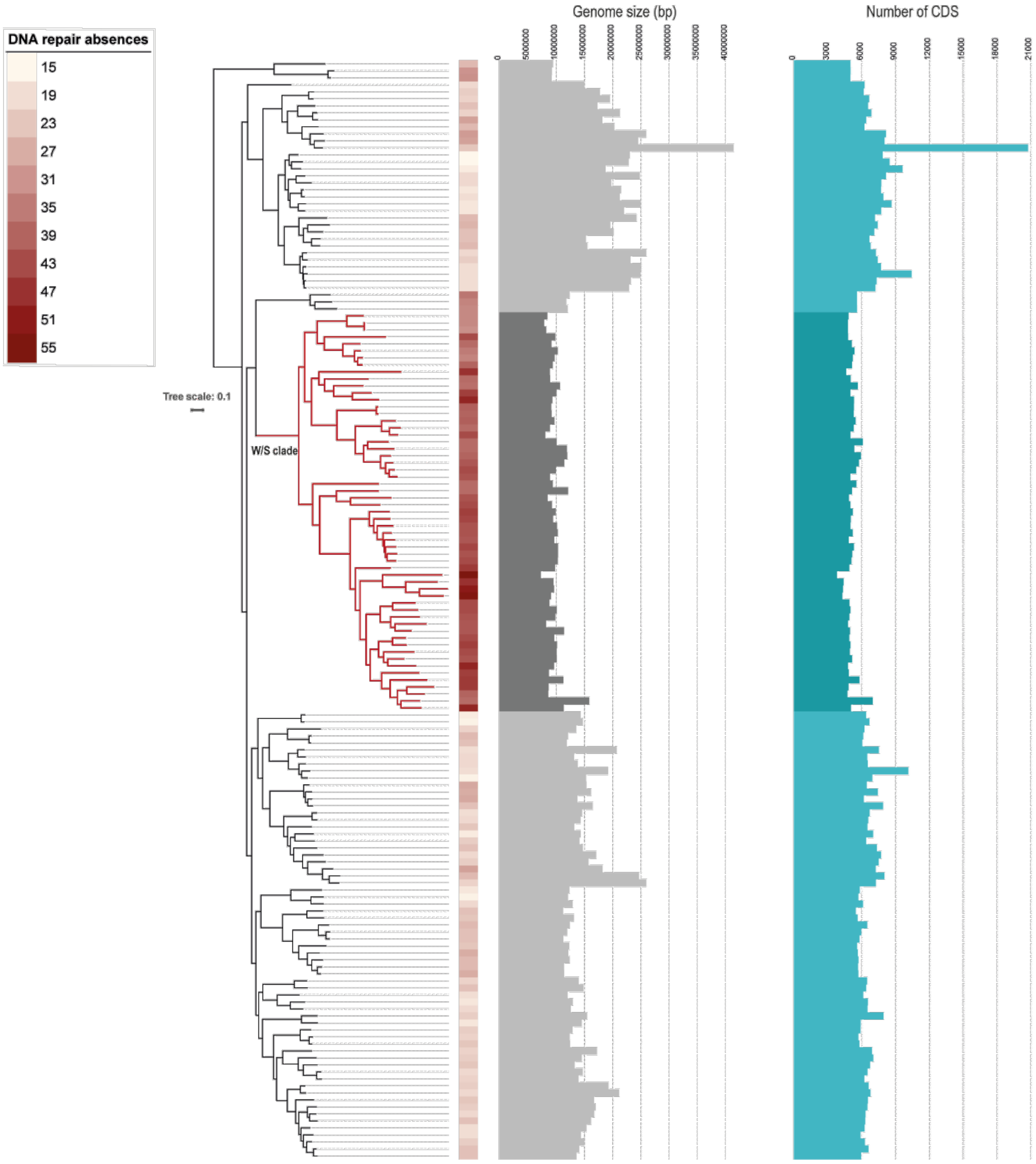


**Figure S9. Species belonging to the W/S clade show evidence of gene loss.** A reduction in genome size and number of coding genes (CDS) can be observed between the W/S clade and closest relatives. The phylogenetic tree, genome size, and CDS count information were obtained from Opulente et al., 2024 (Opulente, et al. 2024).
